# Revisiting the cytogenetics of *Vellozia* Vand.: immunolocalization of KLN1 elucidates the chromosome number for the genus

**DOI:** 10.1101/2024.09.10.612026

**Authors:** Guilherme T. Braz, Lucas B. Riboldi, Maísa S. Pinto, Eliana R. Forni-Martins, Juliana E. C. T. Yassitepe, Ricardo A. Dante, Isabel R. Gerhardt

## Abstract

Chromosome number is the most fundamental trait of a karyotype. Accurate chromosome counting is essential for further analyses including cytogenomics, taxonomic, evolutionary, and genomic studies. Despite its importance, miscounting is common, especially in early publications on species with small and morphologically similar chromosomes. *Vellozia* Vand. is a genus mainly distributed throughout South America belonging to the angiosperm family Velloziaceae, a dominant taxon in the Brazilian “campos rupestres”. Cytogenetic studies within the group have been rare and have shown conflicting chromosome counts, even within the same species. These discrepancies are associated with the presence of a few small chromosome-like structures, which were previously classified as possible satellites. Here, to accurately determine the chromosome number of species belonging to the genus, we used different cytogenomics approaches, including the immunostaining of the KNL1 kinetochore protein combined with chromosome spread preparation using tissue culture-derived samples. Our results revealed 2*n* = 18 chromosomes for all six species studied. This finding suggests that the basic chromosome number for *Vellozia* is *x* = 9 and not *x* = 8, as previously proposed. The immunolocalization of functional centromeres was fundamental for undoubtedly identifying the smaller chromosome pair as real chromosomes and accurately determining the correct chromosome number of these species. This will provide substantial support for further studies, including investigations into karyotype evolution and the generation of reference genomes for the species of the family.

## Introduction

Cytogenetics studies contribute to the analysis of genetic relationships among species and populations, shedding light on the genetic mechanisms driving their diversification (Guerra 2008). The number of chromosomes is the most fundamental karyotype trait and commonly used in taxonomic and evolutionary studies (Guerra 2008). Notably, the generation of reference genomes under current standards requires sequence assembly into chromosome-scale scaffolds that represent the karyotype (Lewin et al. 2018; Whibley et al. 2021; Blaxter et al. 2022; Lawniczak et al. 2022). However, counting chromosomes can be challenging in species with karyotypes composed of numerous, morphologically similar, and small chromosomes (Figueredo et al. 2016). As a consequence, different chromosome numbers have been reported among species or even within the same species, which may not represent real variability but rather result from erroneous counting, especially in earlier studies (Figueredo et al. 2016). Furthermore, errors in taxonomic identification can also be a source of mistakes in chromosomal counting (Guerra 2012). Thus, given the significance of this fundamental karyotype trait, it is crucial to determine the chromosome number of the species under study accurately.

The genus *Vellozia* Vand. belongs to the Velloziaceae family and encompasses about 125 species distributed throughout South America, with only one extending into Panama (WFO 2024). Approximately 118 species are endemic to Brazil, with the vast majority occurring in Minas Gerais State, primarily in the “campos rupestres” (literally, ‘rocky fields’) (Mello-Silva 2005; Alcantara et al. 2018), an open-vegetation, disjunct ecoregion located mostly in the highlands along the Espinhaço Range in eastern Brazil. The “campos rupestres” are considered a biodiversity hotspot due to its high species richness and endemism, with approximately 40% of their angiosperm species being unique to this region (Zappi et al. 2015; Silveira et al. 2016).

The “campos rupestres” are characterized by freely draining and shallow, sandy or rocky soils, high daily temperatures, and an extended dry season occurring through fall and winter (Oliveira et al. 2016; Silveira et al. 2016). The extreme abiotic conditions favor the establishment of species that carry traits related to avoidance of overheating and root specializations allowing an efficient uptake of nutrients, in addition to traits associated with contrasting drought adaptive strategies such as dehydration avoidance and desiccation tolerance (Alcantara et al. 2015; Oliveira et al. 2016; Alcantara et al. 2018). Velloziaceae is an iconic and dominant taxon in the “campos rupestres” (Conceição et al. 2007, 2016; Le Stradic et al. 2015), its species being representative examples of those possessing such traits (Alcantara et al. 2015, 2018). Understanding the mechanisms underlying these traits is a fundamental biological question and can inspire agricultural innovations. This relies on an accurate characterization of these species at different levels, including cytogenomics, where the integration of cytogenetic techniques with genomic data allows researchers to uncover the structural and functional aspects of chromosomes that contribute to the resilience of these plants under stressful conditions.

Chromosomal studies in Velloziaceae are scarce and limited to a few species. The chromosome numbers range from 2*n =* 14 to 2*n =* 48 (Goldblatt and Poston 1988; Melo et al. 1997; Costa et al. 2017; Santos et al. 2018), being the basic number for the family suggested as *x* = 9 (Goldblatt and Poston 1988) or *x =* 8 (Melo et al. 1997). In the genus *Vellozia*, the number of chromosomes ranges from 2*n =* 14 to 2*n =* 18, with its species characterized as diploid (Goldblatt and Poston 1988; Melo et al. 1997; Santos et al. 2018). Importantly, some counts were imprecise, with reports indicating varying numbers within the same sample (2*n =* 14-16 or 2*n =* 14-18) (Melo et al. 1997), revealing difficulties in analysis. These discrepancies are due to a few considerably smaller chromosomes that, in certain preparations, display weak staining. This has led to considering them as possible satellites (Goldblatt and Poston 1988; Melo et al. 1997). Therefore, employing improved chromosome spread preparations combined with more sophisticated cytological analyses is essential for a comprehensive characterization of this group. Although root tips are generally considered the best material for mitotic chromosomal analysis, the root meristem in Velloziaceae species typically contains only a limited number of dividing cells (Melo et al. 1997), making accurate chromosome counts challenging. Therefore, Velloziaceae cytogenetic analyses could benefit from alternative plant samples possessing more actively dividing cells. Additionally, the immunolocalization of kinetochore proteins, like the recently attained for KLN1 (Neumann et al. 2023; Oliveira et al. 2024), may generate more accurate results by marking functional centromeres and enabling the recognition of the smallest chromosomes and/or those with less intense staining.

Given the scarcity of studies and the numerical variability of chromosomes within the genus and even the same species of *Vellozia*, the present study aimed to improve chromosome counting by utilizing short-term callogenesis and centromere immunolocalization; expand the knowledge about the chromosomal number in the genus by determining the chromosome number of six species, among which four previously uncharacterized; and finally contribute to resolving the basic chromosomal number (*x*) in the genus.

## Material and Methods

### Plant material

Mature seeds from *Vellozia intermedia* Seub., *V. nivea* L.B.Sm. & Ayensu, *V. plicata* Mart. (formerly *Nanuza plicata*), *V. aloifolia* Mart., *V. geotegens* L.B.Sm. & Ayensu, and *V. variabilis* Mart. were obtained from the Genomics for Climate Change Research Center (GCCRC) collection.

One population of *V. intermedia* and one of *V. nivea* were assessed for this study, both occupying “campos rupestres” sites (coordinates 20°29’07.2”S, 46°30’49.8”W and 20°29’15.8”S, 46°32’05.3”W, respectively) located on a private property neighboring the Serra da Canastra National Park in Delfinópolis, Minas Gerais state. *V. intermedia* and *V. nivea* individuals were collected, transplanted into pots, and transported to a greenhouse where they were acclimated and cultivated. *V. intermedia*, but not *V. nivea*, individuals flowered in the greenhouse and were manually pollinated at anthesis. Therefore, *V. nivea* individuals had their flowers tagged at anthesis and were allowed to open-pollinate in their “campos rupestres” site of origin. Seeds from both species were harvested from mature fruits no earlier than 30 days after anthesis. Viable seeds of *V. intermedia* were abundantly obtained upon cross-pollination between greenhouse-cultivated individuals.

Seeds of *V. plicata, V. aloifolia, V. geotegens*, and *V. variabilis* were a gift from Dr. Nanuza L. de Menezes (Universidade de São Paulo). *V. plicata* seeds were obtained from individuals cultivated in the garden of the Departamento de Botânica, Instituto de Biociências, Universidade de São Paulo. *V. aloifolia, V. geotegens*, and *V. variabilis* seeds were obtained from individuals growing in “campos rupestres” sites in Diamantina, Minas Gerais (*V. aloifolia* near the BR-367 and MG-220 highways, coordinates 18°16’12.4”S, 43°41’03.6”W; and *V. geotegens* and *V. variabilis* in the Extração/Curralinho district, coordinates 18°16’57.2”S, 43°31’16.3”W).

### Callus generation from mature seeds

Callus induction was performed following published protocols (Deng et al. 2020; Wang et al. 2021), with a few modifications for use in *Vellozia* species. Fifty seeds from each species were surface-sterilized using a two-step process: first, a solution of 70% ethanol and 0.1% Tween 20 (Sigma-Aldrich, Saint Louis, USA) for 30 seconds, followed by 5% sodium hypochlorite and 0.1% Tween 20 for 30 seconds. Post-sterilization, seeds were rinsed twice with distilled deionized water and air-dried before sowing onto the medium.

Murashige and Skoog (MS) basal medium (Sigma-Aldrich) was prepared at a concentration of 4.3 g L^-1^, supplemented with 30 g L^-1^ sucrose, pH adjusted to 5.6, and 10 g L^- 1^ agar added as a solidifying agent. The medium was autoclaved at 120°C for 20 minutes. After autoclaving, an isothiazolinone mixture (Auros Química, São Paulo, Brazil) was added at 0.02% to prevent microbial contamination, along with MS vitamins (0.5 mg L^-1^ thiamine HCl, 0.5 mg L^-1^ pyridoxine HCl, 0.05 mg L^-1^ nicotinic acid, and 2 mg L^-1^ glycine) (Sigma-Aldrich). Plant growth regulators, indole-3-acetic acid (IAA, Sigma-Aldrich) (1 mg L^-1^) and 6-benzylaminopurine (BAP, Sigma-Aldrich) (1 mg L^-1^), were added into the medium for callus induction. Thirty milliliters of the prepared medium were dispensed into 90 × 15 mm Petri dishes.

Ten seeds from each species were placed on the solidified medium in each plate, with five replicates per species. Seed-sown plates were incubated in the dark at 28°C in a growth incubator (Memmert, Schwabach, Germany). Germination started after 20 days, and callus formation was observed on the primary root approximately 10 days post-germination.

### Chromosome spread preparation

Root tips, young leaves, and actively growing callus (20-30 days old) were used to obtain chromosome spreads for *V. intermedia*. For the remaining species, chromosome spreads were obtained only from calli. Root tips and leaves were collected from plants grown in a greenhouse, and calli were obtained as described previously.

All these samples were pre-treated with 0.002M 8-hydroxyquinoline (8HQ) for 5 hours at room temperature (RT). Then, they were fixed in Carnoy solution (3 ethanol: 1 acetic acid) and kept at -20°C until use. Subsequently, they were digested using an enzymatic solution consisting of 4% cellulase R10 (Serva, Heidelberg, Germany), 4% Macerozyme R-10 (Duchefa Biochemie, Haarlem, Netherlands), and 2% cytohelicase (Sigma-Aldrich) for 10 minutes at 37°C. Slides were prepared using the stirring method (Schubert et al. 2001).

### Staining techniques, CMA/DAPI banding, and FISH

We used different strategies, including staining techniques, CMA/DAPI banding, and FISH (Fluorescence *in situ* Hybridization) to initially confirm the chromosome number of *V. intermedia*. For the remaining species, only the immunolocalization of KNL1 was employed, as detailed later. Freshly prepared slides were first stained using 2% acetocarmine in 45% glacial acetic acid solution for an initial investigation.

YOYO-1 (Oxazole Yellow Homodimer; Invitrogen, Waltham, USA), a fluorescent complex enabling ultrasensitive fluorescence detection of nucleic acids, was used to detect the presence of thin chromatin fibers (Rocha et al. 2017). Briefly, 100 μL of YOYO solution consisting of 5 μL YOYO in PBS (1 μL YOYO-1 + 99 μL1X PBS) and 95 μL 1X TRIS-NaCl-blocking buffer (TNB) was applied to the sample and incubated for 30 minutes at 37 °C. Then, they were washed in 1X PBS, dried, and mounted in Vectashield containing DAPI (Vector Laboratories, Newark, USA).

To investigate the occurrence of AT- and GC-rich regions, double staining with CMA/DAPI was applied (Schweizer 1976). Before applying fluorochromes, the slides were left at room temperature (RT) for three days to allow the material to dry. After staining, the slides were stored in the refrigerator to stabilize the fluorescent molecules for at least three days.

The 18S rDNA sequence, one of the three ribosomal DNA genes (18S, 5.8S, and 26S) that compose the 35S rDNA site, was used as a probe in the FISH reaction (Braz et al. 2020). This probe was amplified in a PCR reaction using the genomic DNA from *Solanum pseudocapsicum* L. and the primers NS1 (5’-GTA GTC ATA TGC TTG TCT C-3’) and NS4 (5’-CTT CCG TCA ATT CCT TTA AG-3’) (White et al. 1990). PCR amplification was performed with the GoTaq Green Master Mix (Promega) using the following parameters: 95 ºC for 3 min, 37 cycles at 95 ºC for 1 min, 56 ºC for 1 min, and 72 ºC for 1 min, and 72 ºC for 5 min. The probe was labeled with digoxigenin-11-dUTP (Roche Biochemicals, Burgess Hill, UK) using a nick translation reaction kit (Roche) and detected with an anti-digoxygenin rhodamine-conjugated antibody (Roche). The slides were mounted in Vectashield containing DAPI (Vector Laboratories).

### Chromosome preparation for indirect immunofluorescence

Actively growing calli (20-30 days old) from all six species were pretreated with 8-hydroxyquinoline (8HQ) for 5 hours at room temperature and fixed in ethanol: glacial acetic acid (3:1) for 30 minutes on ice. Subsequently, the callus cells were washed with 1X PBS (Phosphate-Buffered Saline, pH 7.4, Invitrogen), and treated with 4% cellulase Onozuka R10 (Serva), 4% Macerozyme R10 (Duchefa Biochemie), and 2% cytohelicase (Sigma-Aldrich) for 30 minutes at 37 °C. The slides were prepared using the air-drying technique (De Carvalho and Saraiva 1993), omitting the 45% acetic acid treatment. We used at least two callus per slide and prepared three to five slides per species. The slides were stored in 1X PBS at room temperature and subsequently washed in 1X PBS with 0.5% Triton X-100 (Thermo Scientific, Waltham, USA) for 25 minutes.

Rabbit antibodies targeting the kinetochore protein KNL1 (Neumann et al. 2023; Oliveira et al. 2024) were diluted to a concentration of 1:500 in 1X PBS with 0.1% Tween 20. Approximately 200 µL of the diluted antibodies were added to each slide, followed by overnight incubation in a humid chamber at 37 °C. After three washes in 1X PBS, 200 µL of Alexa Fluor 488-conjugated goat anti-rabbit secondary antibody (Invitrogen) (1:500 in 1X PBS with 0.1% Tween 20) were applied to the slides, followed by one-hour incubation in a humid chamber at room temperature. The slides were washed in 1X PBS, and then chromosomes were counterstained with 4,6-diamidino-2-phenylindole (DAPI) in Vectashield antifade solution (Vector Laboratories).

The number of well-spread metaphase chromosomes obtained varied between species and were more than 50 for *V. intermedia*, more than 20 for *V. nivea*, and 2-4 for *V. aloifolia, V. plicata, V. geotegens*, and *V. variabilis*. We were unable to generate immunostaining data for *V. geotegens* and *V. variabilis*. The best chromosome spreads were captured using a DP72 coupled camera attached to an Olympus BX51 epifluorescence microscope and processed with Olympus DP2 BSW (Olympus Corporation, Tokyo, Japan) software. The final image was optimized for brightness and contrast with Adobe Photoshop CS4 (Adobe Systems Incorporated, Mountain View, USA) software.

## Results and Discussion

Chromosome number represents the most fundamental trait of a karyotype, and its precise characterization is vital for cytogenomics research and subsequent studies. Variations in this karyotype feature have been associated with adaptation and speciation, being polyploidy and aneuploidy identified as significant events in the evolution of plant genomes (Guerra 2008; Mayrose and Lysak 2021). Besides, the precise determination of chromosome number is becoming increasingly crucial in today’s “genomics era”. Given the numerous ongoing efforts at assembling reference genomes, it is fundamental to possess knowledge of this trait (Lewin et al. 2018; Whibley et al. 2021; Blaxter et al. 2022; Lawniczak et al. 2022).

Using a short-term tissue culture-based method adapted from others established for seed-derived callogenesis (Deng et al. 2020; Wang et al. 2021), we obtained calli from mature seeds of all six species (Fig. 1A). This yielded a higher number of dividing cells compared to traditional slide preparations from leaf and root tips of *Vellozia* species, from which few suitable cells are typically obtained (Melo et al. 1997), thus facilitating the cytogenetic analysis. Although tissue culture techniques are known to potentially induce somaclonal variation, including the formation of aneuploid cells, these are more often associated with long-term cultures (Roy 1980; Bairu et al. 2011). In contrast, our analysis revealed no variation in chromosome number for *V. intermedia* among cells derived from roots, leaves, or callus. These findings underscore the efficacy of callogenesis as a robust approach for conducting cytogenomics studies in *Vellozia* species.

**Figure 1.**
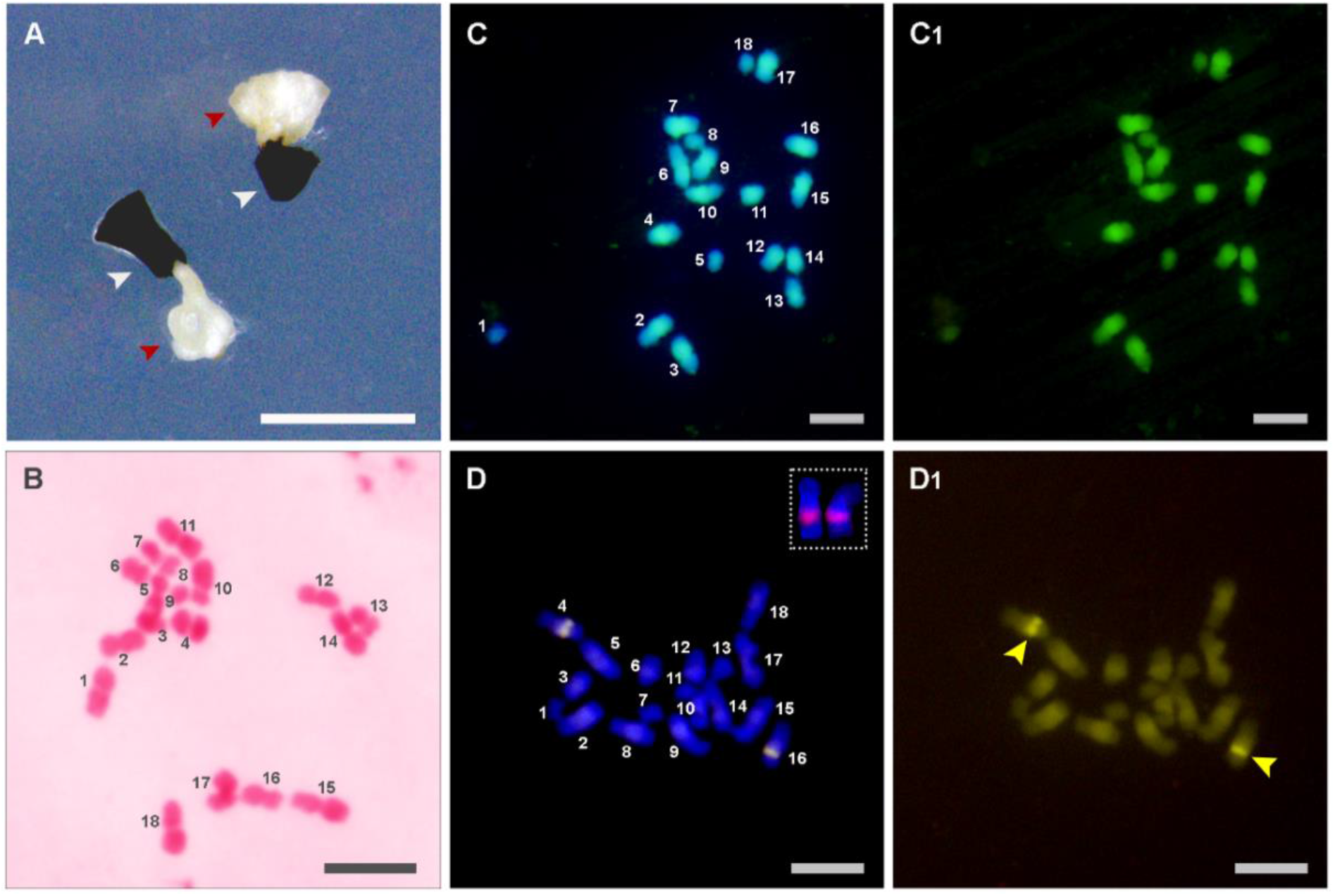
**A** Callus (red arrowhead) induced from mature seeds (white arrowhead) and **B-D1** metaphase chromosomes from *V. intermedia* using different staining techniques. **B** Acetocarmine staining showing 2*n =* 18 chromosomes. **C** Merged DAPI and YOYO staining images. **C1** YOYO staining. **D** CMA/DAPI banding revealing an interstitial pair of CMA^+^ band, probably colocalized to 35S rDNA site (highlighted in the upper right corner). The insert is a higher magnification of a pair of chromosomes containing a 35S rDNA site, obtained from a different metaphase spread. **D1** Pair of chromosomes carrying CMA^+^ band (yellow arrowhead). Scale bars represent 1 mm (A) and 5 µm (remaining images).

The cytogenetics analysis using the acetocarmine staining technique revealed a chromosome count of 2*n =* 18 for *V. intermedia* (Fig. 1B). For over 20 studied *Vellozia* species, the most commonly reported chromosome numbers are 2*n =* 14, 2*n =* 16 or 2*n =* 14-16 chromosomes (Supplementary Table 1). Exceptions include *V. glabra*, which has been reported to have chromosome counts ranging from 2*n* = 14-18 (Melo et al. 1997), and *V. plicata*, with a count of 2*n* = 18 (Santos et al. 2018), despite previous descriptions indicating a chromosome number of 2*n* = 16 (Melo et al. 1997).

All these uncertainties about chromosome counts across *Vellozia* species arise from the observation of the shadowy chromatic bodies and chromosome-like structures, which were interpreted as satellites and not classified as chromosomes (Goldblatt and Poston 1988; Melo et al. 1997). Therefore, we used different cytogenomics approaches to confirm the nature of these chromatin bodies. In *Lolium* L. species, YOYO staining was successfully applied to detect fragile sites associated with 35S rDNA, revealing chromatin fiber connecting acentric chromosomal fragments (satellites) and centromere-containing chromosomes (Rocha et al. 2017). Using the same approach, we were unable to visualize any chromatin fibers linking the potential chromosomal fragments, such as the smaller pairs of chromosome-like structures, to centromere-containing chromosomes (Fig. 1C, C1). Therefore, this result suggests that our current resolution may not have been sufficient to visualize highly decondensed chromatin fibers, or alternatively, it suggests that the observed smaller chromosome-like structures may indeed represent real chromosomes.

The CMA banding technique revealed a single pair of CMA^+^ bands located interstitially on the long arm of a chromosome pair (Fig. 1D, D1). Additionally, FISH using 18S rDNA as a probe also identified a single pair of this site, likely colocalized with the CMA^+^ band (Fig. 1D). Thus, these results suggest that, in contrast to observations in *Lolium* (Rocha et al. 2017), the 35S rDNA in our study does not induce fragile sites or the formation of satellites, supporting our finding that the smallest chromosome-like structures indeed represent chromosomes.

Finally, to conclusively confirm the chromosome number of *V. intermedia*, we immunolocalized functional centromeres using an universal antibody that targets seed plant KLN1 kinetochore proteins (Neumann et al. 2023; Oliveira et al. 2024). This approach, consistent with our previous findings (Fig. 1B), revealed eighteen individual immuno-signals that confirms the presence of 2*n =* 18 monocentric chromosomes in this species (Fig. 2).

**Figure 2.**
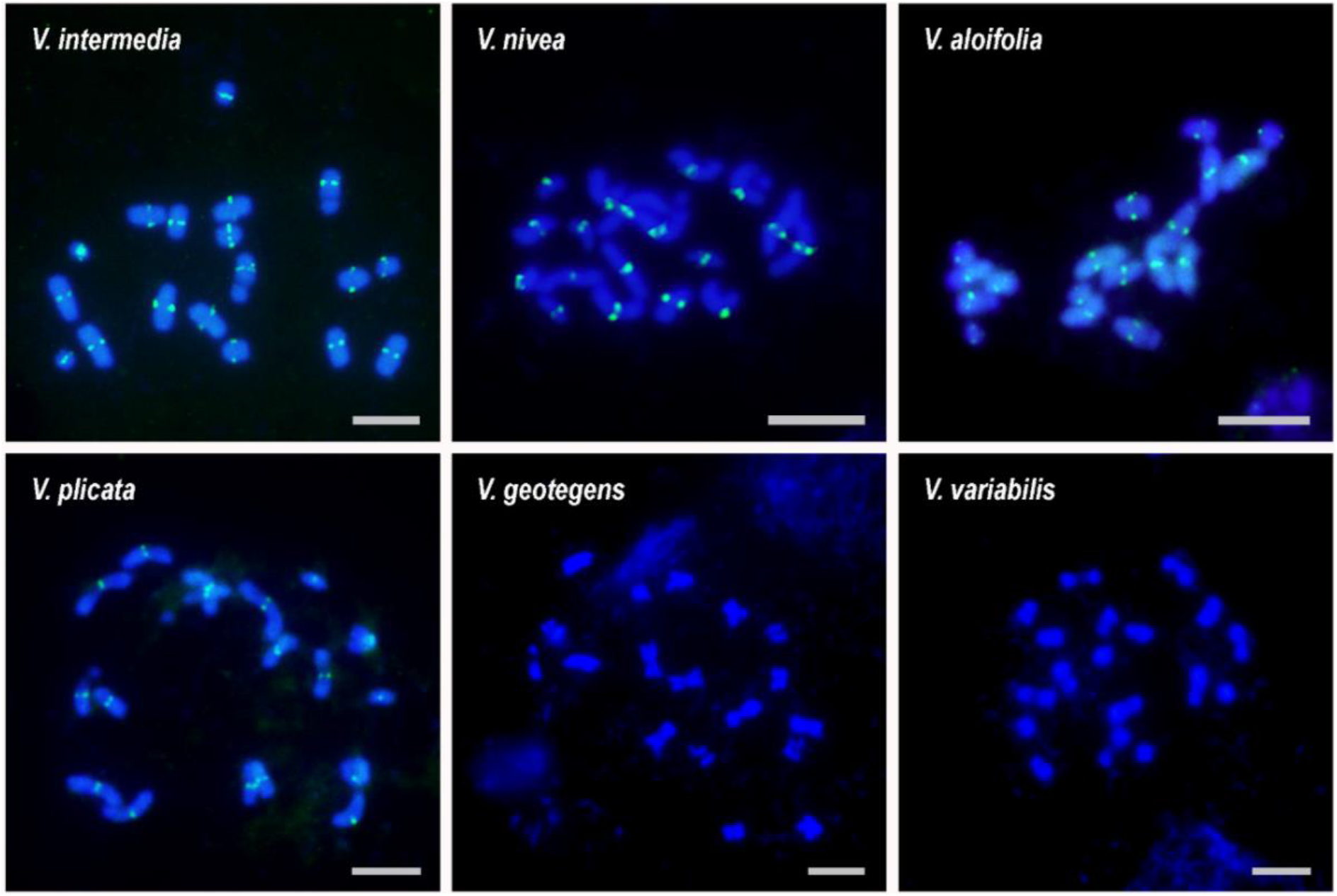
Immunolocalization of the KLN1 protein on metaphase chromosomes obtained from calli of *V. intermedia, V. nivea, V. aloifolia*, and *V. plicata* revealing 18 functional centromeres and confirming the chromosome count of 2*n =* 18. We were unable to generate immunostaining results for *V. geotegens* and *V. variabilis*. However, DAPI staining indicated the same chromosome number for these species. Scale bars represent 5 µm.

Furthermore, we investigated whether the chromosome count 2*n =* 18, observed in *V. intermedia*, is also present in other *Vellozia* species. We successfully obtained many dividing cells in callus from all five remaining species. However, we achieved a high number of metaphases with well-spread chromosomes only in the *V. nivea* samples, indicating that adjustments in the protocol are necessary for the other species. Nevertheless, the 2*n =* 18 count was in fact observed in all five other species (Fig. 2). This includes *V. geotegens* and *V. variabilis*, for which we were unable to perform immunostaining (Fig. 2). The chromosome number of 2*n =* 18 revealed for *V. nivea* contrasts with the previously described 2*n =* 16 (Melo et al. 1997). Thus, our findings establish a new chromosome count for this species while confirming the count of 2*n =* 18 chromosomes for *V. plicata* (Santos et al. 2018). This is the first report of the chromosome numbers for *V. aloifolia, V. geotegens*, and *V. variabilis*, in addition to *V. intermedia*.

The application of the immunostaining technique to identify functional centromeres was fundamental to determine the correct chromosome number for *Vellozia*. In addition, the consistent number in different species excludes the hypothesis that the accessions studied here represent cytotypes, harbor B-chromosomes, or exhibit aneuploidy resulting from callogenesis. Importantly, because our analysis encompasses *Vellozia* species that are distantly related phylogenetically (Alcantara et al. 2018), the resulting chromosome numbers are representative of the whole genus. Therefore, our results suggest that the chromosome number for all *Vellozia* species is likely 2*n =* 18, indicating *x =* 9 as the basic number for the genus and not *x =* 8 as previously proposed (Goldblatt and Poston 1988; Melo et al. 1997).

Through a combination of callogenesis and centromere immunolocalization, we overcame challenges resulting from the limited number of dividing cells in meristems and the small size of the chromosomes, and consequently contributed to resolving a decades-long controversy regarding the chromosome number in the *Vellozia* genus. This will considerably support further studies on these dominant and iconic species of the Brazilian “campos rupestres”, including investigations into karyotype evolution in the genus and Velloziaceae in general, and the generation of chromosome-scale reference genomes.

## Supplementary Information

The online version contains supplementary material available.

## Acknowledgments

This work was supported by grants 2016/23218-0 and 2021/00280-0 from Fundação de Amparo à Pesquisa do Estado de São Paulo (FAPESP). G.T.B. (2021/00224-2) and M.S.P. (2022/04929-3) received post-doctoral fellowships from FAPESP, and L.B.R. (173550/2023-1) a post-doctoral fellowship from Conselho Nacional de Desenvolvimento Científico e Tecnológico (CNPq). We would like to express our gratitude to Dr. Nanuza L. de Menezes for *Vellozia* seeds, Dr. Jiri Macas and Dr. Ludmila Oliveira (Institute of Plant Molecular Biology, Czech Academy of Sciences) for the donation of the KNL1 antibody, and Recanto Ecológico Vale do Céu (Delfinópolis, MG) for granting access to the “campos rupestres” sites.

## Authors’ contributions

R.A.D and I.R.G. conceived the project and collected individuals from the “campos rupestres” sites. All authors contributed to the study design. G.T.B. collected cytological data. L.B.R., M.S.P. and J.E.C.T.Y. developed the tissue culture method and produced calli. All authors contributed to the data analysis. G.T.B. and E.R.F.M. drafted the manuscript. All authors wrote, reviewed and approved the submission of the manuscript. G.T.B and L.B.R. contributed equally to this study.

## Data availability

All data included in this study are available upon request by contacting the corresponding authors.

