## Supplementary Table 1 for "Revisiting the cytogenetics of *Vellozia* Vand.: immunolocalization of KLN1 elucidates the chromosome number for the genus"

**Supplementary Table 1** Published chromosome numbers (*n* and 2*n*) of Velloziaceae species, including the current valid names, the original names used in the corresponding publications, and the respective bibliographic references (Ref.).

| **Current valid name** | **Original name used in the publication** | ***n*** | **2*n*** | **Ref.** |
| --- | --- | --- | --- | --- |
| *Ac. bracteata* P.C.Kao | *Ac. bracteata* P.C.Kao | - | 38 | K |
| - | *Aylthonia* sp. nova | - | 34 | M |
| - | *B*. aff. *albiflora* L.B.Smith | 17 | - | GP |
| *B. coronata* Ravenna | *B. coronata* P. Ravenna | 17 | - | GP |
| *B. fanniae* (N.L.Menezes) Mello-Silva | *P. fanniei* N. Menezes | - | 34 | M |
| *B. globata* Goethard & Henrard | *B. globata* Goethard & Henrard | 17 | - | GP |
| *B. graminifolia* L.B.Sm. | *A. graminifolia* (L.B.Smith) N.Menezes, comb. nov. ined. | - | 34 | M |
| *B. inclinata* Goethart & Henrard | *P. inclinata* (Goethard & Henrard) N. Menezes | - | 34 | M |
| *B. longiscapa* Goethart & Henrard | *P. longiscarpa* (Goethard & Henrard) N.Menezes | - | 34 | M |
| *B. nuda* L.B.Sm. & Ayensu | *P. nuda* (L.B.Smith & Ayensu) N. Menezes, comb. nov. ined. | - | 34 | M |
| *B. paranaensis* L.B.Sm. | *B. paranaensis* L.B. Smith [= A. paranaensis (L.B.Smith) N.Menezes] | 17 | - | GP |
| *B. pulverulenta* L.B.Sm. &Ayensu | *A. pulverulenta* (L.B.Smith) N.Menezes | - | 34 | M |
| *B. pungens* (N. L. Menezes & Semir) Mello-Silva | *Bu. pungens* (N.Menezes & Semir | - | 34 | M |
| *B. purpurea* Hook. | *P. purpurea* (Hooker) Raf. | - | 34 | M |
| *B. riparia* (N. L. Menezes & Mello-Silva) Mello-Silva | *P. riparia* N.Menezes & Mello-Silva | - | 34 | M |
| *B. rogieri* T. Moore & Aires | *P. rogieri* (Moore & Aires) N.Menezes, comb. nov. ined. | - | 34 | M |
| *B. spiralis* L.B.Sm. & Ayensu | *Bu. spiralis* (L.B.Smith & Ayensu) N.Menezes & Semir | - | 34 | M |
| *V. abietina* Mart. | *X. abietina* (Mart.) Sprengel | - | 14 | M |
| *V. alata* L.B.Sm. | *V. alata* L.B.Smith | 7 | - | GP |
| *V. albiflora* Pohl | *V. crassicaulis* Mart. ex J.A, & J.H. Schult. (= V. albiflora Pohl | - | 14-16 | M |
| *V. aloifolia* Mart. | - | - | 18 | * |
| *V. bahiana* L.B.Sm. & Ayensu | *V. bahiana* L.B.Smith & Ayensu | - | 14 | M |
| *V. bahiana* L.B.Sm. & Ayensu | *V. bahiana* L.B.Smith & Ayensu | 8 | - | GP |
| - | *V*. aff. *candida* Mikan | - | 14 | M |
| *V. candida* J.C.Mikan | *V. candida* Mikan | - | 14-16 | M |
| *V. caruncularis* Mart. ex Seub. | *V. caruncularis* Mart. ex Seubert | 7 | - | GP |
| *V. compacta* Mart. ex Schult. & Schult.f. | *V. compacta* Mart. ex Seubert | 8 | - | GP |
| - | *V.* aff. *declinans* Goethard & Henrard | - | 14 | M |
| *V. geotegens* L.B.Sm. & Ayensu | - | - | 18 | * |
| *V. giuliettiae* N. Menezes & Mello-Silva | *X. giuliettiae* N.L.Menezes & Semir, sp. nov. ined. | - | 14 | M |
| *V. glabra* J.C.Mikan | *V. glabra* Mikan | - | 14-18 | M |
| *V. grao-mogulensis* L.B.Sm. | *V. grao-mogulensis* L.B.Smith | - | 16 | M |
| *V. hirsuta* Goethart & Henrard | *V. hirsuta* Goethard & Henrard | 7 | - | GP |
| *V. hirsuta* Goethart & Henrard | *V. riedeliana* Goethard & Henrard | 7 | - | GP |
| *V. intermedia* Seub. | - | - | 18 | * |
| *V. minima* Pohl | *X. minima* (Pohl) Baker | - | 14 | M |
| *V. nanuzae* L.B.Sm. & Ayensu | *V. nanuzae* L.B.Smith & Ayensu | - | 16 | M |
| *V. nivea* L.B.Sm. & Ayensu | *V. nivea* L.B.Smith & Ayensu | - | 16 | M |
| *V. nivea* L.B.Sm. & Ayensu | - | - | 18 | * |
| - | *V.* aff. *patens* L.B.Smith & Ayensu | - | 14 | M |
| V. patens L.B Sm. & Ayensu | *V. patens* L.B.Smith & Ayensu | - | 16 | M |
| *V. plicata* Mart | *N. plicata* (Mart.) L.B.Smith & Ayensu | - | 16 | M |
| *V. plicata* Mart. | *V. plicata* Mart. | - | 18 | S |
| *V. plicata* Mart. | - | - | 18 | * |
| *V. pterocarpa* L.B.Sm. & Ayensu | *V. pterocarpa* L.B.Smith & Ayensu | 8 | - | GP |
| *V. pusilla* Pohl | *V. pusilla* L.B.Smith & Ayensu | - | 16 | M |
| *V. sellovii* Seub. | *X. sellovyi* (Seubert) Baker | - | 14 | M |
| *V. squamata* Pohl | *V. flavicans* Mart. ex Schult. | - | 16 | M |
| *V. tubiflora* (A. Rich.) Kunth | *V. tubiflora* (A.Rich.) Kunth | 7 | - | GP |
| *V. variabilis* Mart. ex Schult. & Schult. f. | - | - | 18 | * |
| *V. variegata* Goethart & Henrard | *V. variegata* Goethard & Henrard | - | 14-16 | M |
| - | *Vellozia* sp. nova | - | 16 | M |
| *X. dasylirioides* Baker | *X. dasylirioides* Baker var. pectinata (Baker) H.Perr | - | 48 | M |
| *X. elegans* Baker | *X. elegans* (Balf.) Baker [= *Talbopiopsis elegans* (Hook f.) L.B.Smith | - | 48 | M |
| *X. elegans* Baker | *X. elegans* (Balf.) Baker [= *Talbopiopsis elegans* (Hook f.) L.B.Smith | 24 | - | GP |
| *X. elegans* Baker | *X. elegans* (Balf.) Baker [= *Talbopiopsis elegans* (Hook f.) L.B.Smith | - | ~ 48, 52 | SS |
| *X. humilis* T. Durand & Schinz | *X. humilis* (Baker) Dur. & Schinz | 24 | - | GP |
| *X. humilis* T. Durand & Schinz | *X. humilis* Th.Dur. & Schinz | - | 48 | H |
| *X. retinervis* Baker | *X. retinervis* Baker | 24 | - | GP |
| *X. schlechteri* Baker | *X. viscosa* Baker | 24 | 48 | C |

Genera: *Ac* = *Acanthochlamys*, *A* = *Aylthonia*, *B* = *Barbacenia*, *Bu* = *Burlemarxia*, *N* = *Nanuza*, *P* = *Pleurostima*, *V* = *Vellozia*, and *X* = *Xerophyta*. References: C = Costa et al. 2017, GP = Goldblatt & Poston (1988), H =Hanson (2001), K = Kao et al. (1993), M = Melo et al. (1997), S = Santos et al. (2018), SS = Svensson-Stenar (1925), *this work.
